# ADAR1 orchestrates the ZBP1-mediated PANoptosis and heart transplant rejection

**DOI:** 10.1101/2024.09.05.611429

**Authors:** Haitao Lu, Jifu Jiang, Xuyan Huang, Aaron Haig, Lakshman Gunaratman, Anthony M. Jevnikar, Zhu-Xu Zhang

## Abstract

**Background:** PANoptosis is an integrated form of cell death that combines features of pyroptosis, apoptosis, and necroptosis and is regulated by a complex network of signaling proteins. The roles of ADAR1 (adenosine deaminase acting on RNA 1) and RIPK1 (receptor-interacting serine/threonine-protein kinase 1) in orchestrating the ZBP1 (Z-DNA binding protein 1)-RIPK3 complex to mediate PANoptosis is not fully understood, particularly in the context of heart transplantation.

**Objective:** This study investigated how ADAR1 and RIPK1 coordinate the activation of the ZBP1-RIPK3 complex to mediate PANoptosis and its implications in mouse heart transplantation.

**Methods:** Using both in vitro and in vivo models, we analyzed the interactions between ADAR1, RIPK1, ZBP1, and RIPK3. We employed western blotting, and siRNA to elucidate the dynamics of these interactions. Additionally, we assessed the impact of ZBP1 on mouse heart transplantation outcomes.

**Results:** Our studies revealed that ADAR1 regulates the activation of the ZBP1-RIPK3 complex for PANoptosis. The interaction of ADAR1 with ZBP1 protected against Z-DNA-induced cell death by limiting activations of ZBP1 and RIPK3. In mouse heart transplantation study, we found that ZBP1 and its ligand Z-DNA/Z-RNA were significantly increased in the graft post-transplantation. Furthermore, ZBP1 deficiency in the heart graft inhibited cardiac PANoptosis, attenuated acute graft injury, and induced long-term graft survival.

**Conclusion:** This study elucidates the role of ADAR1 in ZBP1-mediated PANoptosis. Inhibition of ZBP1 can prevent heart graft injury and rejection. Understanding these mechanisms provides valuable insights into the regulation of cell death and may inform the development of novel therapeutic strategies to improve transplant outcomes.

## Introduction

Heart transplantation is a critical treatment for end-stage heart failure, offering a life-saving option for patients with severe cardiac conditions. Despite advancements in surgical techniques and immune-suppressive therapies, transplant-related injuries remain a significant challenge, affecting graft survival and patient outcomes. Previous studies have identified various mechanisms contributing to transplant injuries, including ischemia-reperfusion injury and inflammatory responses.^1^ Recently, the role of programmed cell death pathways, such as PANoptosis, has gained attention for its involvement in organ injuries. PANoptosis has been identified as an integrated form of cell death that combines features of apoptosis, necroptosis, and pyroptosis, reflecting a complex interplay of these pathways ^2^.

Circulating cell-free DNAs (cfDNAs) are fragments of DNA released into the bloodstream during cell damage or stress and have emerged as important mediators of inflammation responses ^3^. cfDNAs can act as damage-associated molecular patterns (DAMPs), influencing the activation and regulation of various cell death pathways. cfDNAs have been used as valuable monitors for graft rejection in different stages of transplantation^4^. The increase of Z-form DNA (Z-DNA) among 3-type DNAs (A-, B-, and Z-form) correlates closely with the activation of cell death pathways ^5^. Subsequently, Z-DNA or Z-RNA can be recognized by nucleic sensors, especially, Z-DNA-binding protein 1 (ZBP1) and adenosine deaminase acting on RNA 1 (ADAR1).

Sensing Z-nucleic acid by the ZBP1 Zα domain stimulates ZBP1 to interact with receptor-interacting protein kinase 3 (RIPK3) through RIP homotypic interaction motif (RHIM) homotypic interaction, thereby activating inflammatory cell death ^6^. ZBP1 and RIPK3 are central to the regulation of PANoptosis in the context of virus or fungi infection ^7, 8^. However, it has not been studied whether ZBP1 promotes Z-DNA-induced cell death in transplant injury.

ADAR1-p150 contains a Zα domain, which recognizes endogenous nucleic acids ^9^. Besides ADAR1, ZBP1 is the only other mammalian protein containing Zα domains^10^. ADAR1 suppresses ZBP1-mediated cell death via the Zα interaction.^11^ ADAR1 deficiency increased cell death^9^. However, the mechanism of how ADAR1 regulates the ZBP1-RIPK3 complex is still unclear.

This study is to investigate how ADAR1 regulates the ZBP1-RIPK3 complex to mediate PANoptosis in cardiac microvascular endothelial cells. We demonstrated that ADAR1 orchestrates the activation of the ZBP1-RIPK3 complex in the induction of PANoptosis. Elucidating the role of ADAR1 in PANoptosis has significant implications for our understanding of cell death regulation. Furthermore, we found that ZBP1 deficiency in the heart graft attenuated PANoptosis, graft injury, and rejection, implying a potential therapeutic strategy in transplantation.

## Materials and Methods

### Mice

C57BL/6 (B6) mice and BALB/c mice (Charles River Lab, Bar Harbor, ME, USA) were maintained in the animal facility of the University of Western Ontario. The ZBP1 gene knockout (ZBP1^-/-^) mice were kindly gifted from Dr. Kanneganti TD ^11^, St. Jude Children’s Research Hospital, Memphis, USA, with the permission of Dr. Akira S, Osaka University^12^. All animal experimental procedures comply with the rules and guidelines of the Institutional Animal Care and Use Committee and are approved by the Animal Care Committee of Western University (AUP-2023-135).

### Cell culture and cell death assay

Human cardiac microvascular endothelial cells (HVECs) were purchased from Lonza and were immortalized through transfection of SV40-T-antigen (MiliporeSigma). Cells were grown in DMEM medium supplemented with fetal bovine serum. Cell death was induced as we previously demonstrated (REF). Briefly, TNFα (10 ng/mL; PeproTech, NJ, USA), Smac-mimetic BV6 (2 μM; ApexBio Tech. Houston, TX, USA) and caspase-8 inhibitor IETD (30 μM; ApexBio) were added for 12 hours when cells reached the maximum death. Then the supernatant was used for DNA extraction (K182002, ThermoFisher). Then cfDNAs were used for induction of PANoptosis.

HVECs were seeded in 96-well plates and were pretreated with cold 0.1M CaCl2 for 5 minutes and washed by Opti-MEM (Gibco, 31985070). cfDNAs (final concentration: 5 ng/μl) were transfected into cells with Lipofectamine 300 (Invitrogen, L3000001). Cell death was detected by propidium Iodide (PI) or SYTOXR Green (100 nM, Thermo Fisher Scientific, S7020) and analyzed by Cytation 5 (Agilent, CA, United States). Cells seeded in 6-well plates were used for PCR or Western Blotting.

To further confirm cell death, ATP levels in HVECs were measured by the ATP Bioluminescence Kit (G9241, Promega, USA). In addition, cell death leads to the release of Lactate dehydrogenase which was measured by CyQUNT kit (Invitrogen, C20301).

### dsDNA and Z-DNA ELISA

DNA ELISA assays were described previously^13^. Briefly, Immulon 2HB plates (ThermoFisher) were coated with 100 μL/well of DNAs from supernatants overnight at 4 °C in 1X SSC (150 mM NaCl, 15 mM sodium citrate, pH 7.0). Control wells were 1X SSC alone. The plates were washed 3 times with 1X Phosphate Buffered Saline (PBS) and block buffer (2% bovine serum albumin (BSA), 0.05% Tween-20 in 1X PBS) for 2 hours. After blocking, plates were washed 3 times with PBS. The plates were then incubated for 1 hour with anti-Z-DNA antibody (Absolute Antibody, Z22) or mouse anti-ds-DNA antibody (Thermofisher, # 606-430). All antibodies were diluted with Tris dilution buffer (0.1% BSA, 0.05% Tween 20 in 50 mM Tris, pH 7.4). After incubation with primary antibodies, the ELISA plates were washed 3 times with 1X PBS and incubated with 100 μL/well of the horseradish peroxidase (HRP)-conjugated anti-mouse IgG (γ chain specific) at 1:1000. The plates were washed 3 times with PBS and then incubated for 30 min in the dark with 100 μL/well of a substrate mixture (0.015% 3, 3’, 5, 5’-tetramethylbenzidine dihydrochloride, 0.01% H_2_O_2_ in 0.1 M citrate buffer, pH=4). Then, 100 μL/well of 2 M H_2_SO_4_ was added to terminate the reaction. The absorbance was measured at 450 nm (Multi-plate spectrophotometer, Molecular Devices, San Jose, CA, USA).

Brominated-poly (dG: dC) (Br-poly(dG: dC) was added as the positive control to transform Z-DNA ^14, 15^. Briefly, the poly (dG: dC) (Invivogen) is dissolved in bromine-saturated solution (Fisher Scientific) in 20 mM sodium citrate (pH 7.2), 1 mM EDTA, and 4 M NaCl. The reaction proceeded at room temperature for 10 min. One milligram of Br-poly(dG-dC) has an A260 of 14.4. The solution was kept at room temperature to facilitate the B-to Z-DNA transition.

### RNA silencing

HVECs were seeded at 50–60% confluence. HVECs were transfected with the siRNA working solution in serum-free Opti-MEM (Thermo Fisher Scientific) and Lipofectamine 3000 (Thermo Fisher Scientific) according to the manufactory protocol. siADAR1 (SMARTPool, Dharmacon RNAi Tech), siRIPK3 (Thermofisher), and scrambled siRNA (Dharmacon RNAi Tech) were used.

HVECs were transfected with siRNA for 24 hours for PCR analysis and 48 hours for Western blotting. Based on the PCR and Western blotting results, only the HVECs with over 90% knockdown rate would be selected for this study.

shZBP1-1 CCAAGTCCTCTACCGAATGAA; shZBP1-2 GCACAATCCAATCAACATGAT within the pLKO.1 vector (MiliporeSigma, TRCN0000123050) were transfected into HVECs using Lipofectamine 3000 (Invitrogen, L3000001). After transfection, puromycin was used to perform cell selection with increased concentration (0.5-10 μg/ml) for 7 days. PCR and western blotting were used to confirm the knockdown efficiency.

### Real time PCR

Total RNA extraction was used by Qiagen RNeasy Mini Kit (Qiagen,74104). PCR was performed using PowerTracker QPCR Mix (Thermo Fisher Scientific). The primers were: ADAR1: 5’-ATGCTCCTCCTTTCCAGGTC-3’ and 5’-TGGTCAGAGCATTGAAGCAC-3’. ZBP1: 5’-TGGTCATCGCCCAAGCACTG-3’ and 5’-GGCGGTAAATCGTCCATGCT-3’. AIM2: 5’-AGGCTGCTACAGAAGTCTGTCC-3’ and 5’-TCAGCACCGTGACAACAAGTG G3’. cGAS: 5’-TTCCACGAGGAAATCCGCTGAG-3’ and 5’ CAGCAGGGCTTCCTGGTTTT TC-3’. IFI16: 5’-GATGCCTCCATCAACACCAAGC-3’ and 5’-CTGTTGCGTTCAGCACCATC AC-3’. TMEM17: 5’-CCTGAGTCTCAGAACAACTGCC-3’ and 5’-GGTCTTCAAGCTGCCCACAGTA-3’. Caspase 1: 5’-GCTGAGGTTGACATCACAGGCA-3’ and 5’-TGCTGTCAGA GGTCTTGTGCTC-3’. IL-1: 5’-AGTAGCAACCAACGGGAAGG-3’ and 5’-TTCCTCTGAGT CATTGGCGA-3’. GSDMD: 5’-ATGAGGTGCCTCCACAACTTCC-3’ and 5’-CCAGTTCC TTGGAGATGGTCTC-3’. β-actin: 5’-CCAGCCTTCCTTCCTGGGTA-3’ and 5’-CTAGAACATTGCGGTGCA-3’. β-actin was used as endogenous control.

### Western Blotting

HVECs were lysed with lysis buffer (BioRad) followed by high-speed centrifugation (13000 rpm,10 min) to remove debris. Protein concentration was measured by Bradford Dye Protein Assay (ThermoFisher Scientific) before electrophoresis and electrophoretic transfer.

PVDF membranes were blocked by 5% skim milk and incubated with the following primary antibodies respectively: anti-caspase-1 (CST,#2225), anti-cleaved-caspase-1 (CST,# 4199),anti-caspase-3 (CST, #9662), anti-caspase-4/5 (CST, #42264), anti-cleaved-caspase-4/5 (Invitrogen, # PA5-39873),anti-cleaved caspase-3 (CST, #9661), anti-caspase-7 (CST, #9492), anti-cleaved caspase-7 (CST, #9491), anti-caspase-8 (CST, #4927), anti-cleaved caspase-8 (CST, #8592), anti-caspase-9 (CST, # 9502), anti-cleaved caspase-9 (CST, # 9505), Rat anti-Caspase-11 (CST, #14340), anti-cleaved caspase-11(Santacruz, sc-56038) anti-p-RIPK3 (CST, #91702S), anti-RIPK3 (CST, # 10188), anti-p-RIPK1 (CST, # 65746), anti-RIPK1 (CST, #3493)anti-p-MLKL (CST, # 18640), anti-MLKL (CST,# 14993), anti-GSDMD (CST,# 39754), anti-cleaved-GSDMD (CST, human#36425); anti-cleaved-GSDMD (CST, mouse# 10137), anti-GSDME (CST,#19453), anti-cleaved-GSDME(CST,# 55879), anti-ADAR1 (CST, #14175), anti-ZBP1 (CST, human #60968),anti-ZBP1 (CST, mouse #84056), TNFR1(CST, #3736), TLR3(CST, #6961), Phospho-NF-κB p65 (Ser536, CST,#3033) anti-GAPDH (CST,# #2118). The dilution of each primary antibody was initially 1:1000 in 2% BSA buffer.

Membranes were then washed and incubated with the appropriate horseradish peroxidase (HRP)–conjugated secondary antibodies: anti-rabbit (Abcam, ab288151) or anti-mouse (Abcam, ab97261) for 1 hour. Proteins were visualized by using Immobilon Western Chemiluminescent HRP Substrate (Millipore, WBKLS0500).

### Heterotopic Cardiac Transplantation

Donor hearts from B6 or ZBP1^-/-^ mice were transplanted into allogeneic BALB/c mice according to the approved protocol ^16^. Briefly, mice were anesthetized by ketamine/xylazine, and hearts were procured through a butterfly thoracic incision and the median sternotomy. To induce transplantation IRI, the donor’s heart was perfused with Ringer’s buffer and stored at 37°C for 1 hour and at 4 °C for 4 hours. The graft was anastomosed to the recipient’s abdominal aorta and inferior vena cava using 11-0 sutures (Ethicon, Piscataway, NJ, USA). The recipients were kept on an inhaled isoflurane/oxygen mixture (MilliporeSigma). The abdominal wall and skin were closed with a 5-0 suture (Ethicon, Piscataway, NJ, USA). Anti-CD154 (0.25mg/mouse,740874, BD Biosciences, San Jose, CA, USA) was injected intraperitoneally right after transplant surgery.

For assessing acute injury, the graft recipients were euthanatized three days after for histology analysis. In addition, graft survival was monitored by abdominal palpation. Cessation or a significant drop of cardiac pulsation was considered graft rejection.

### Histology

Grafts were perfused with saline and cut transversely, then fixed with 5% formalin. Paraffin sections were used for hematoxylin and eosin (H&E) staining, the TdT-mediated dUTP nick-ended labeling (TUNEL), and immunohistochemistry (IHC).

Artery damage, infarction, PMN, and leukocyte infiltration were evaluated by a pathologist in a blind manner. Injury was scored on a scale of 0–5 (0: no change, 1: 0–10% change,2: 10–25% change, 3: 25–50% change, 4: 50–75% change, 5: >75% changes).

Intra-graft death in tissue sections was detected by the TUNEL method (Millipore, Sigma). The number of TUNEL-positive cells was quantified by Image J in a double-blinded fashion.

For lymphocyte detection, tissue sections were detected by anti-CD3 (Abcam, ab243874) and anti-CD45 (Abcam, ab10558). For confirmation of ZBP1-mediated PANoptosis, following antibodies were used for IHC: anti-Z-DNA (Absolute Antibody, Z22), anti-dsDNA (Thermo Fisher, mouse# 606-430), anti-ZBP1(CST, mouse #84056), anti-cleaved GSDMD (CST, mouse #10137), anti-cleaved GSDME (CST, mouse #38821), anti-cleaved caspase 3 (CST, #9661), anti-p-MLKL (CST, # 18640).

## Statistical Analyses

Data analysis was performed by the student’s t-test or 1-way ANOVA with Tukey’s post-hoc corrections test. The Mantel-Cox log-rank test was used to determine graft survival differences. Differences were considered significant when p-value ≤ 0.05.

## Result

### 1. Z-DNA in cfDNAs Promotes ZBP1 Mediated PANoptosis

cfDNAs have been reported as post-transplantation markers in clinics. ^17-19^ Z-DNA concentration increases during DNA fragmentation ^20, 21^. We detected Z-DNA levels in cfDNA samples using Z-DNA antibody and ELISA. Cell death was induced as we previously described ^16, 22, 23^. cfDNAs were extracted from supernatants and analyzed by ELISA. A significant increase of Z-DNA in cfDNA samples was detected (Figure 1A).

**Figure 1.**
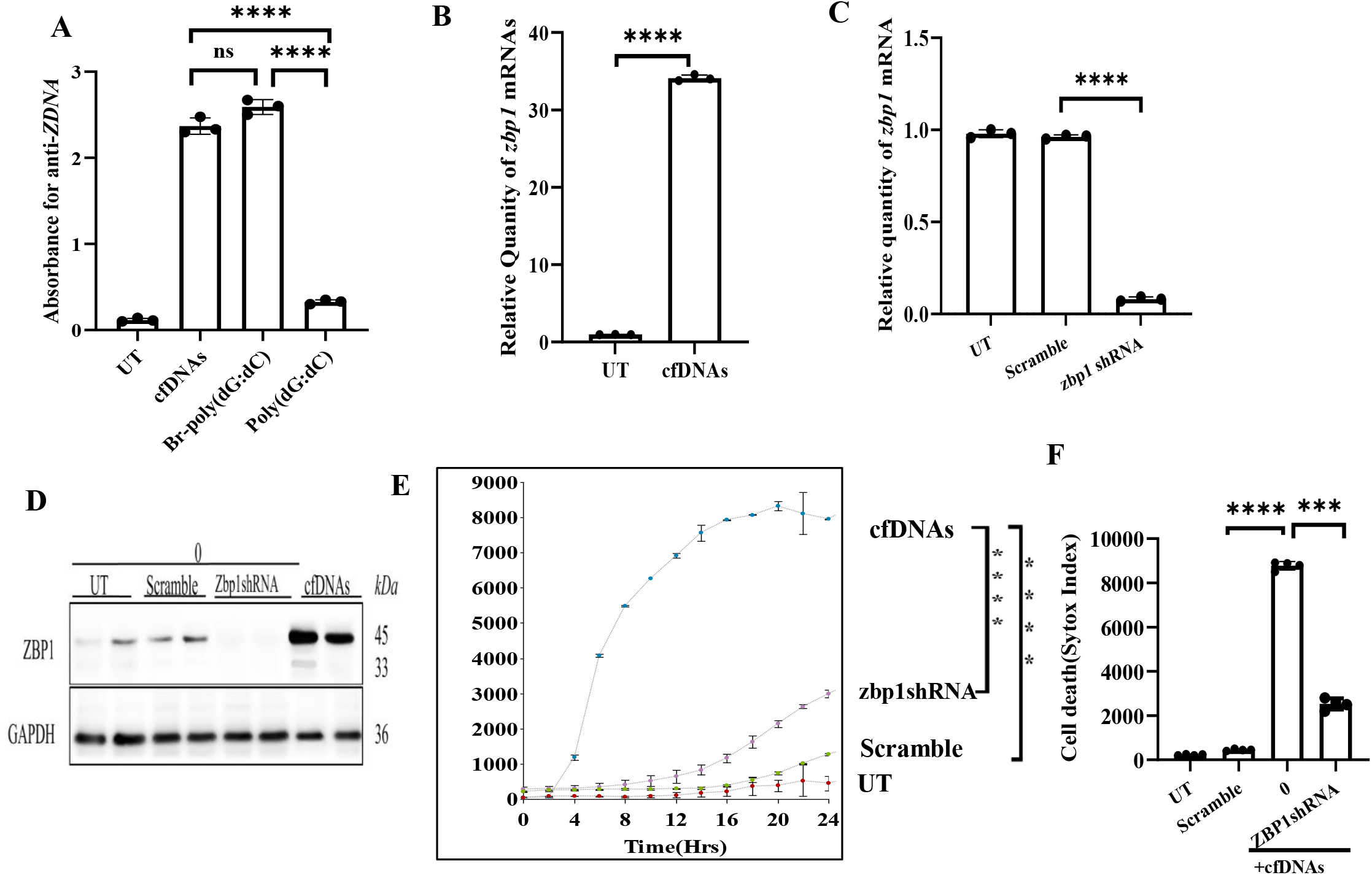
Increased Z-DNA in cfDNAs promotes ZBP1-mediated cell death. (A) Z-DNA ELISA. Cells (1× 10^4^/well) were seeded in triplicates in a 96-well plate and treated with TNFα (T, 20 ng/mL), Smac mimetic BV6 (S, 2 μM) and caspase-8 inhibitor z-IETD (I, 30 μM) to induce necroptosis for 12 hours. The supernatant was collected and analyzed by ELISA with anti-Z-DNA. The brominated (Br)-poly (dG: dC) treated group was the positive control of Z-DNA, and poly (dG: dC) was the negative control. Data are shown as mean± SD of triplicates. (B) Expression of ZBP1 was quantified by real-time PCR after cfDNAs (5ng/ul) treatment for 3 hrs. β-actin was used as an endogenous control for mRNA expression. Data are shown as mean ± SD of three independent experiments. (C) Expression of ZBP1 was quantified by real-time PCR after ZBP1 shRNA silencing. Data are shown as mean ± SD of three independent experiments. (D) Western blot analysis of ZBP1 in cells treated with control, shRNA, or cfDNAs alone. (E&F) Cell death assay. Cells (1 × 10^4^/well) were seeded in quadruplicates in a 96-well plate and treated with cfDNA (5 ng/ul) premixed with Lipofectamine3000. Cell death was detected by PI uptake from 0 to 24 h and monitored by Cytation 5. PI uptake was quantified at 24 h. Data are shown as mean± SD of quadruplicates and representative of three independent experiments. (H) Colocalization of ZBP1 and Z-DNA. Cells were treated with cfDNAs for 6 hours and stained with anti-ZBP1-PE and anti-Z-DNA-FITC as detailed in the Methods. The overlay (Orange) of PE (red) and FITC (green) indicates the binding of ZBP1 and Z-DNA. *** p,0.001, **** p < 0.0001. t-test

Next, we analyzed ZBP1 in response to cfDNA. ZBP1 mRNA level was increased significantly in cfDNA-treated cells, and this increase was inhibited by ZBP1 shRNA (Figure 1B&C). Western blot analysis confirmed changes of ZBP1 (Figure 1D).

In our cell death assays, cfDNAs (5 ng/μl) promoted cell death compared to the UT group (Figure 1E&F). ZBP1 inhibition by shRNA significantly prevented cell death (Figure 1E&F). In summary, those results supported the notion that Z-DNAs promote cell death via ZBP1.

Next, we determined if cfDNA treatment could induce PANoptosis, a form of cell death that integrates features of pyroptosis, apoptosis, and necroptosis. Disulfiram was added to inhibit the activation of GSDMD/E, Z-VAD to inhibit caspases, and NSA to inhibit MLKL. Interestingly, each of those inhibitors partially reduced cell death (Figure 2A-B), which was also confirmed by analysis of ATP levels (Figure 2C).

**Figure 2.**
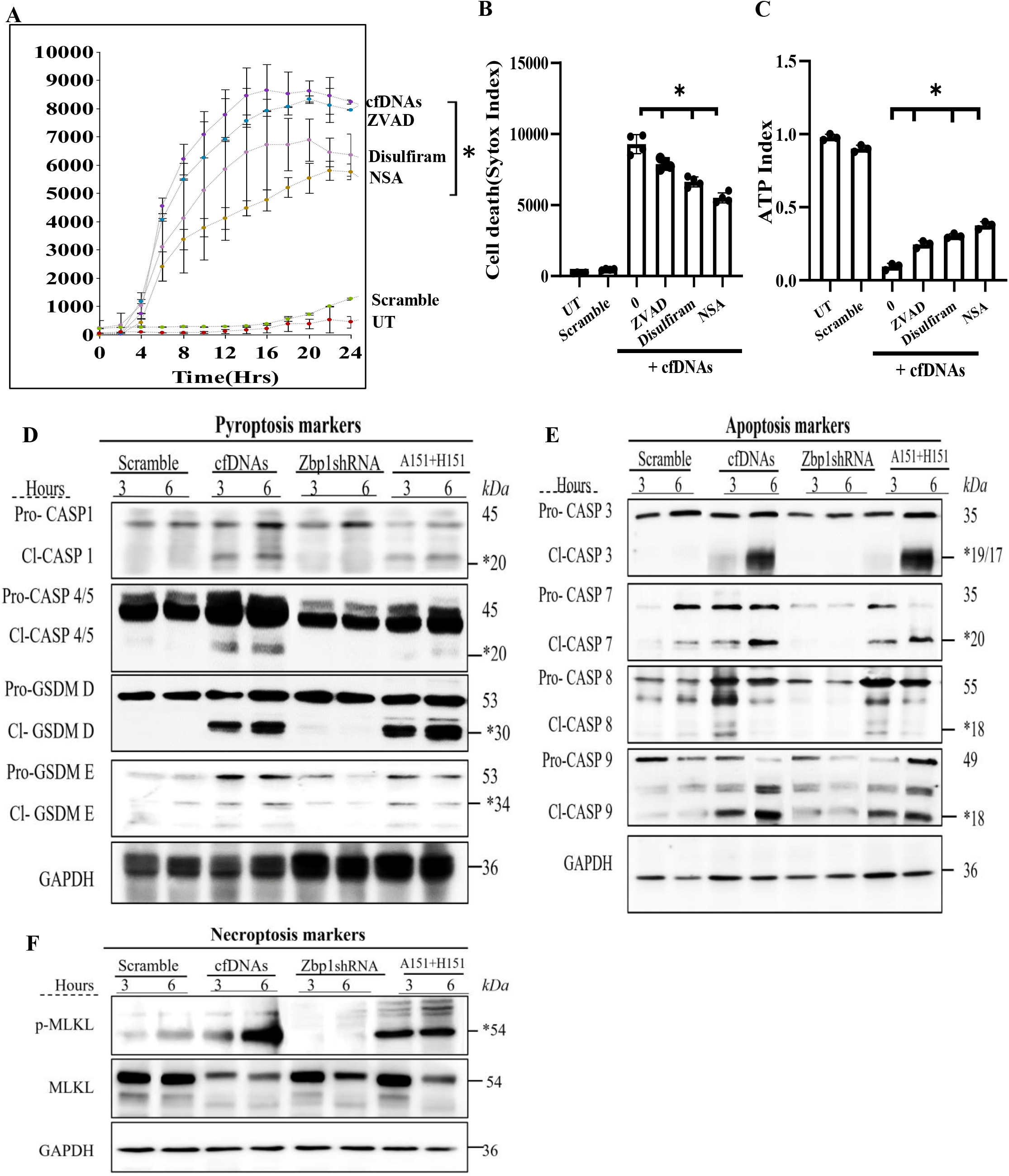
ZBP1-mediated PANoptosis. (A&B). Cells were treated with cfDNAs (5ng/ul) together with inhibitor to pyroptosis (Disulfiram, GSDMD/E inhibitor, 20 uM), apoptosis (ZVAD, pan-caspases inhibitor, 30 uM), necroptosis (NSA, p-MLKL inhibitor, 10 uM) respectively. Cell death was detected by PI uptake from 0 to 24 h and monitored by Cytation 5. PI uptake was quantified at 24 h. Data are shown as mean± SD of quadruplicates and representative of three independent experiments. *p < 0.05. t-test. (C) Cell death was confirmed by detecting ATP level. (D-F) Western blot analysis of pyroptosis, apoptosis, and necroptosis markers. Cells were treated with cfDNA and ZBP1 shRNA, A151 (inhibitor of AIM2 and cGAS, 10uM) plus H151 (STING inhibitor, 2uM) for 3 hours or 6 hours. Scramble siRNA was used as a control. Expressions of (caspase-1/4/5, GSDMD, GSDME, caspase-3/7/8/9, and pMLKL were detected by antibodies. “*” represents the cleaved form of the molecule.

Next, we detected PANoptosis molecules after cells were treated with cfDNAs for 6 hours. cfDNAs treatment induced cleaved GSDMD (P30) (Figure 2D), indicating pyroptosis. GSDMD can be activated by cleaved caspase-1 (p20) or caspase-4/5 (p20) which were increased as well (Figure 2D)^24^. Another member of the gasdermin family, GSDME, was also increased. In addition, increases in caspases 3, -7, -8, and -9 indicated apoptosis (Figure 2E). pMLKL increase indicates the occurrence of necroptosis (Figure 2F). Taken together, these results suggested that cfDNA treatment induces PANoptosis.

To confirm the role of ZBP1 in PANoptosis, we used shRNA to silence ZBP1. ZBP1 knockdown significantly inhibited cleaved caspase-3, GSDMD, and pMLKL, collectively PANoptosis (Figure 2D-F). However, other studies showed that AIM2 or cGAS-STING participates in PANoptosis^21,25^. To clarify if this is the case, we treated cells with AIM2 or STING inhibitor A151 (25 uM) and STING inhibitor H151 (50 uM). The combination of A151 and H151 did not rescue cell death (Figure 3C&D). Western blot (Figure 2D-F) showed that PANoptosis molecules were not inhibited by A151 and H151. Taken together, these data indicate that ZBP1, not dsDNA sensors, plays a central role in mediating PANoptosis.

**Figure 3.**
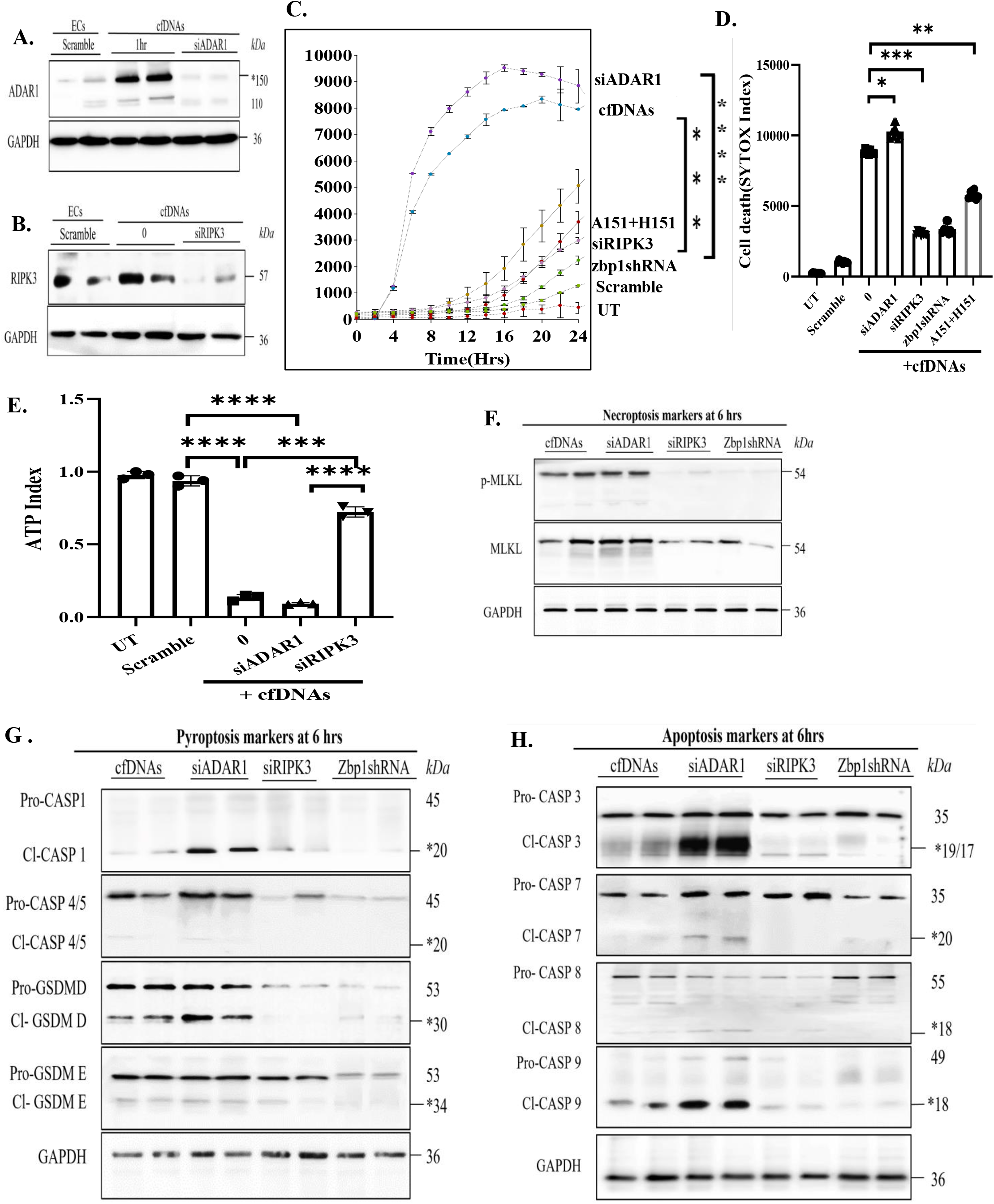
The loss of ADAR1 exacerbates ZBP1-mediated PANoptosis. (A&B) Cells were treated with ADAR1 siRNA or RIPK3 siRNA. Scramble siRNA was used as a control. 24 hours later, cell death was induced by cfDNAs as detailed in the Methods, and cells were collected at 3 or 6 hours for Western blot analysis of ADAR1 and RIPK3. (C&D). Cells were treated with ADAR1 siRNA, RIPK3 siRNA, and ZBP1 shRNA, respectively. Scramble siRNA was used as a control. 24 hours later, cell death was induced as above. Cell death was detected by PI uptake and monitored by Cytation 5. PI uptake was quantified at 24 h. Data are shown as mean± SD of quadruplicates and representative of three independent experiments. *p < 0.05, **p < 0.01, ***p < 0.001, **** p < 0.0001 t-test. (E) The ATP level was measured. (ns (not significant), *p < 0.05, **p < 0.01, ***p < 0.001, **** p < 0.0001 t-test. (F-H) Western blot analysis of pyroptosis, apoptosis, and necroptosis markers in cells treated with cfDNAs (5 ng/ul) for 6 hours. (“*” represents the cleaved form of the molecule).

### 2. ADAR1 and RIPK3 regulate ZBP1-Mediated PANoptosis

ADAR1-p150 isoform and ZBP1 are the only two proteins with the Zα domain. In addition, Z-DNA/RNA stimulates ZBP1 or ADAR1 by recognizing the Zα domain ^20, 26^. Therefore, ADAR1 may regulate ZBP1’s function in responding to Z-DNA/RNA. To clarify their relationship, ADAR1 was silenced by siRNA as confirmed by PCR and Western blot (Figure 3A and Supplement Figure 2A). ADAR1 knockdown exacerbated cell death (cell death index 10500± 660 vs 8600± 540, Figure 3C&D). In addition, the ATP assay confirmed that the loss of ADRA promoted cell death (Figure 3E). These results suggest that ADAR1 may inhibit ZBP1-mediated cell death.

Next, we determine the role of RIPK3 in ZBP1-mediated PANoptosis. It is reported that Z-DNA promotes the interaction between ZBP1 and RIPK3 through the RHIM domain, resulting in PANoptosis^9, 27^. The complex ZBP1-RIPK3 recruits caspases for apoptosis and MLKL for necroptosis. In addition, ZBP1-RIPK3 can recruit caspase-8 and caspase-6 to form a cell death signaling scaffold which stimulates nucleotide-binding domain and leucine-rich-repeat family pyrin domain containing 3 (NLRP3) inflammasome, leading to pyroptosis ^28^.

Interestingly, RIPK3 silencing (Figure 3B) significantly reduced cell death (Figure 3C&D). Cell death inhibition was confirmed as RIPK3 silencing restored ATP levels (Figure 3E). Taken together, RIPK3 participated in Z-DNA-induced cell death.

Next, we determined if RIPK3 is the downstream molecule of ZBP1 to form the PANoptosome. First, cleaved GSDMD (p30) was increased by ADAR1 knockdown but decreased by RIPK3 or ZBP1 knockdown (Figure 3F). Activating caspase-1 and caspase-4/5 in ADAR1 knockdown enabled the cleavage of GSDMD (Figure 3F). ADAR1 knockdown activated caspases 3, 7, 8, and 9, while RIPK3 or ZBP1 knockdown attenuated the activation of caspases (Figure 3G). The necroptosis marker, pMLKL, was promoted by ADAR1 knockdown but inhibited by RIPK3 or ZBP1 knockdown (Figure 3H). Taken together, the absence of ADAR1 exacerbates PANoptosis triggered by ZBP1-RIPK3.

### 3. Transplant injury promotes Z-DNA, ZBP1, ADAR1, and PANoptosis in the graft

To validate our in vitro findings, we studied PANoptosis in mouse heart transplantation. Donor hearts from wild-type B6 mice or ZBP1^-/-^ mice were subjected to ischemic storage as detailed in the Methods before being transplanted into allogeneic BALB/c mice. Immunohistochemistry (IHC) analysis showed significantly increased levels of Z-DNA/RNA in heart grafts 3 days post-transplantation (Figure 4A&B). This is consistent with the increase of Z-DNA in sera post-transplantation as shown in Figure 1B. We observed a significantly elevated level of ADAR1 in ZBP1^-/-^ grafts compared with wild-type grafts, as confirmed by IHC (Figure 4C&D) and western blot (Figure 5A&C). Whereas, ZBP1 expression increased in wild-type grafts post-transplantation, as confirmed by IHC (Figure 4E&F) and western blot (Figure 5A&B). These data indicate a regulatory interaction between ADAR1 and ZBP1.

**Figure 4.**
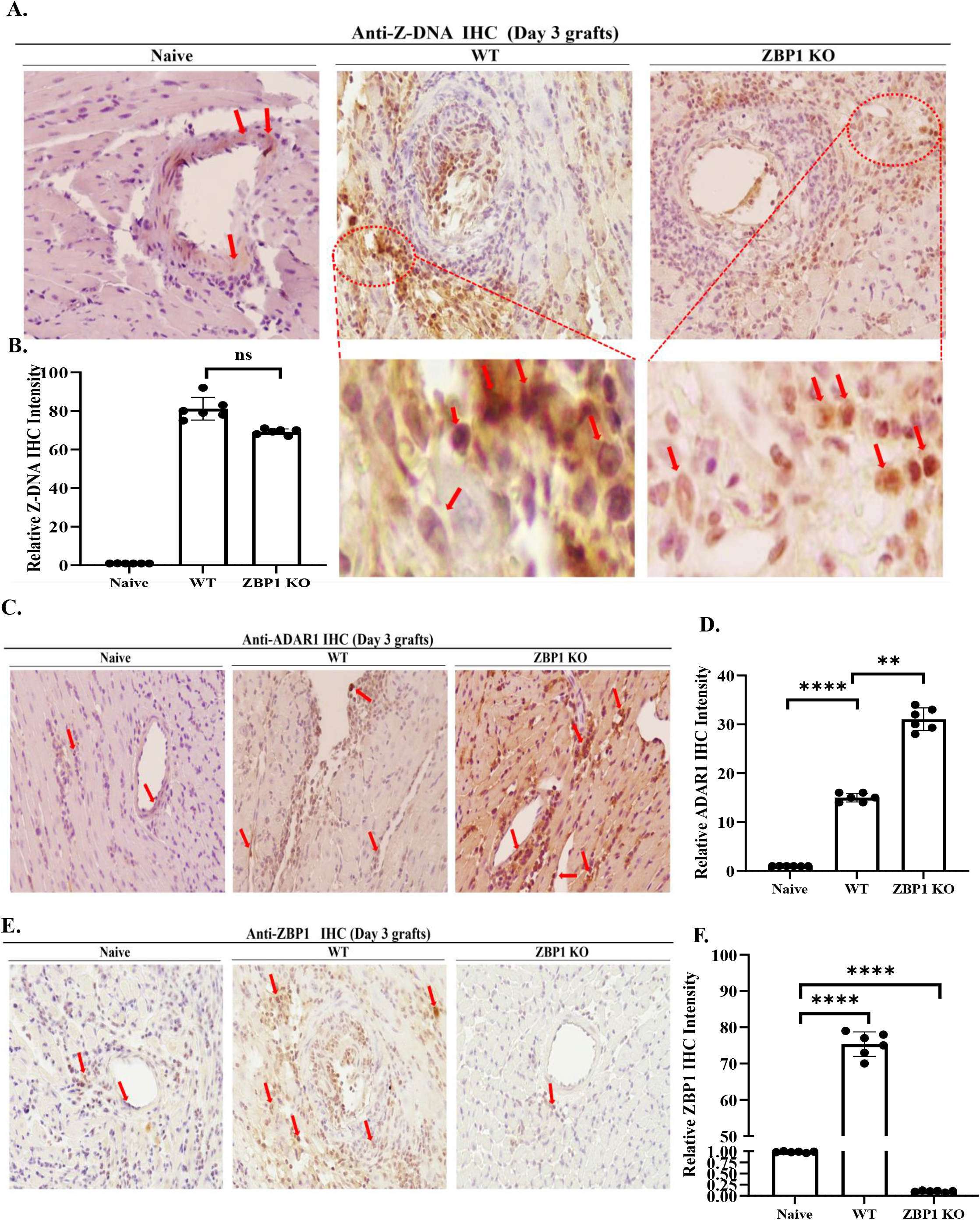
Transplant injury promotes Z-DNA, ZBP1, and ADAR1 expressions in the graft. (A&B) B6- or ZBP1^-/-^-to-BALB/c heart transplantation are described in Materials and Methods. Grafts (n = 6) were collected after 3 days for anti-Z-DNA immunohistochemistry (IHC). Red arrows indicate positive areas (brown color). Images were taken under 200 times magnification. Each graft was automatically counted in six connected random areas by Image J and averaged in a double-blinded fashion. Scores were averaged as mean ± SD of 6 grafts. (C&D) anti-ADAR1 IHC. Red arrows indicate positive areas (brown color). Images were taken under 200 times magnification. Each graft was automatically counted in six connected random areas by Image J and averaged in a double-blinded fashion. ns (not significant), **p < 0.01, ***p < 0.001, t-test. (E&F) anti-ZBP1 IHC. Red arrows indicate positive areas (brown color). Images were taken under 200 times magnification. Each graft was automatically counted in six connected random areas by Image J and averaged in a double-blinded fashion. ns (not significant), ***p < 0.001, ****p < 0.0001, t-test.

**Figure 5.**
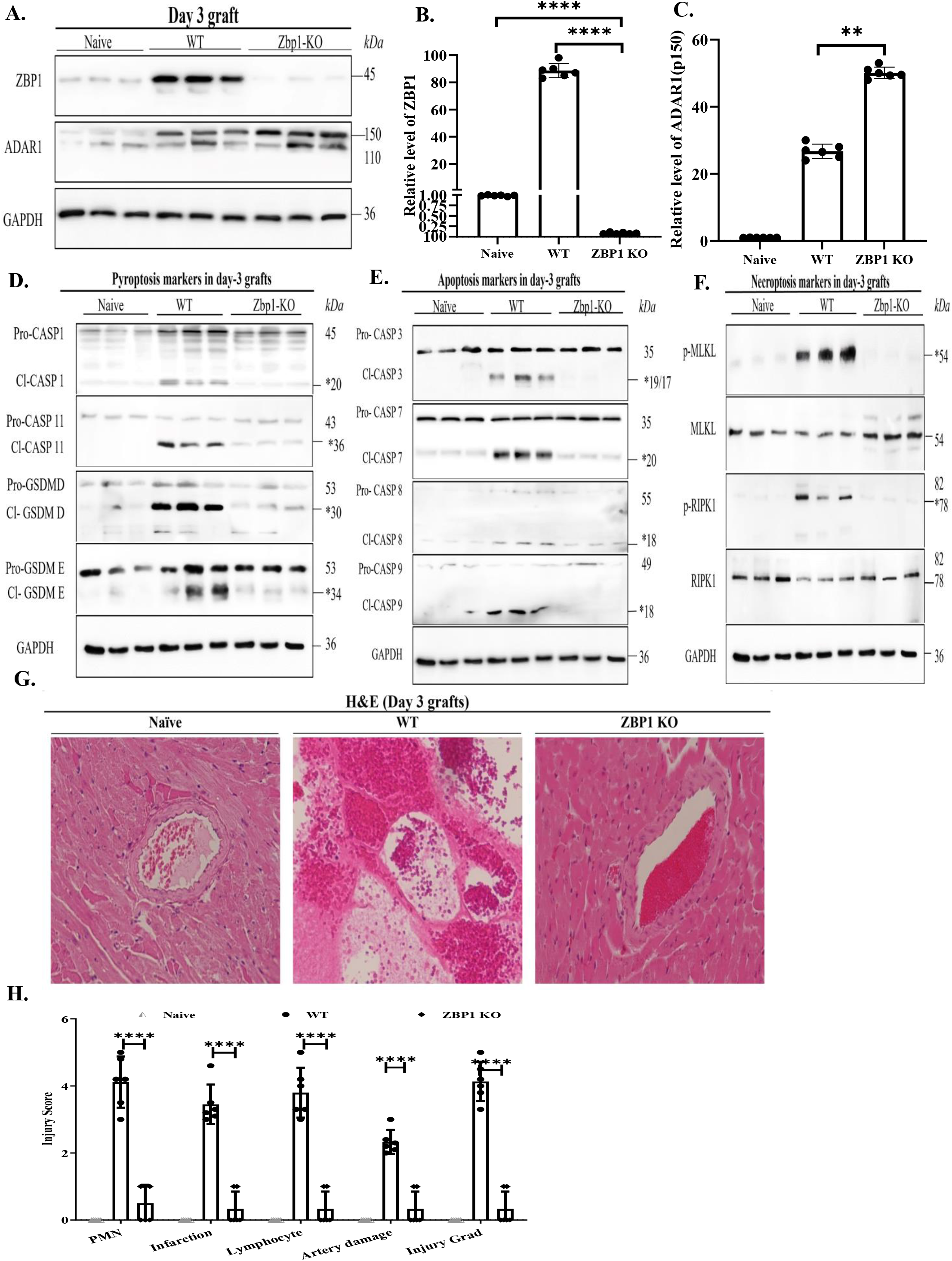
Transplant injury promotes PANoptosis in the graft. (A-C) Western blot analysis of ZBP1 and ADAR1. Data are shown as mean± SD of three grafts. (D-F) Western blot analysis of pyroptosis (caspase-1 and -11, GASFM-D and -E), apoptosis (caspase -3,-7, -8, -9), and necroptosis (MLKL and RIPK1). The results are representative of three different grafts. (G-H) Grafts (n = 6/group) were collected after 3 days of transplantation for H&E staining. Images were taken under 200 times magnification. Graft injuries were quantified blindly by a pathologist. Scores were averaged as mean ± SD. *p < 0.05, **p < 0.01, ***p < 0.001, ****p < 0.0001, t-test.

To understand the impact of ADAR1 and ZBP1 interaction, we analyzed markers of PANoptosis. Western blot analysis showed elevated levels of cleaved-GSDMD and GSDME, activated caspases 1-11, and pMLKL (Figure 5D-F). These data confirmed that the transplant injury is associated with PANopotosis.

### 4. ZBP1 deficiency in the graft attenuated acute graft injury and induced long-time transplant survival

Our above studies indicated that the transplant injury is associated with PANopotosis, and ZBP1 deficiency prevented PANoptosis. Therefore, ZBP1 deficiency in the graft may prevent graft injury and rejection. Next, we analyzed tissue damage post-transplantation. Grafts were collected 3 days after transplantation and stained by H&E. The tissue sections were scored by a pathologist in a double-blinded fashion. There was a significantly reduced injury in ZBP1^-/-^ grafts compared to WT controls (Figure 5G&H). ZBP1^-/-^ grafts have less polymorphonuclear leukocyte (PMN) infiltration (0.5 ± 0.5 vs. 4.1 ± 0.9 in control, n = 6, p = 0.00019), lymphocyte infiltration (0.3 ± 0.6 vs. 3.8 ± 0.7 in control, n = 6, p = 0.00048), infarction (0.3 ± 0.5 vs. 3.5 ± 0.8 in control, n = 6, p = 0.00017), artery damage (0.3 ± 0.7 vs. 2.3 ± 0.9 in control, n = 6, p = 0.048), and overall injury (0.4 ± 0.5 vs. 4.2 ± 0.8, n = 6, p = 0.00015).

Next, we analyzed cell death in the grafts. TUNEL showed that the cell death area was significantly reduced in the ZBP1^-/-^ heart graft (Figure 6A&B). Similarly, a decrease in graft lymphocyte infiltration was confirmed by IHC using anti-CD3 and anti-CD45 (Figure 6C-F).

**Figure 6.**
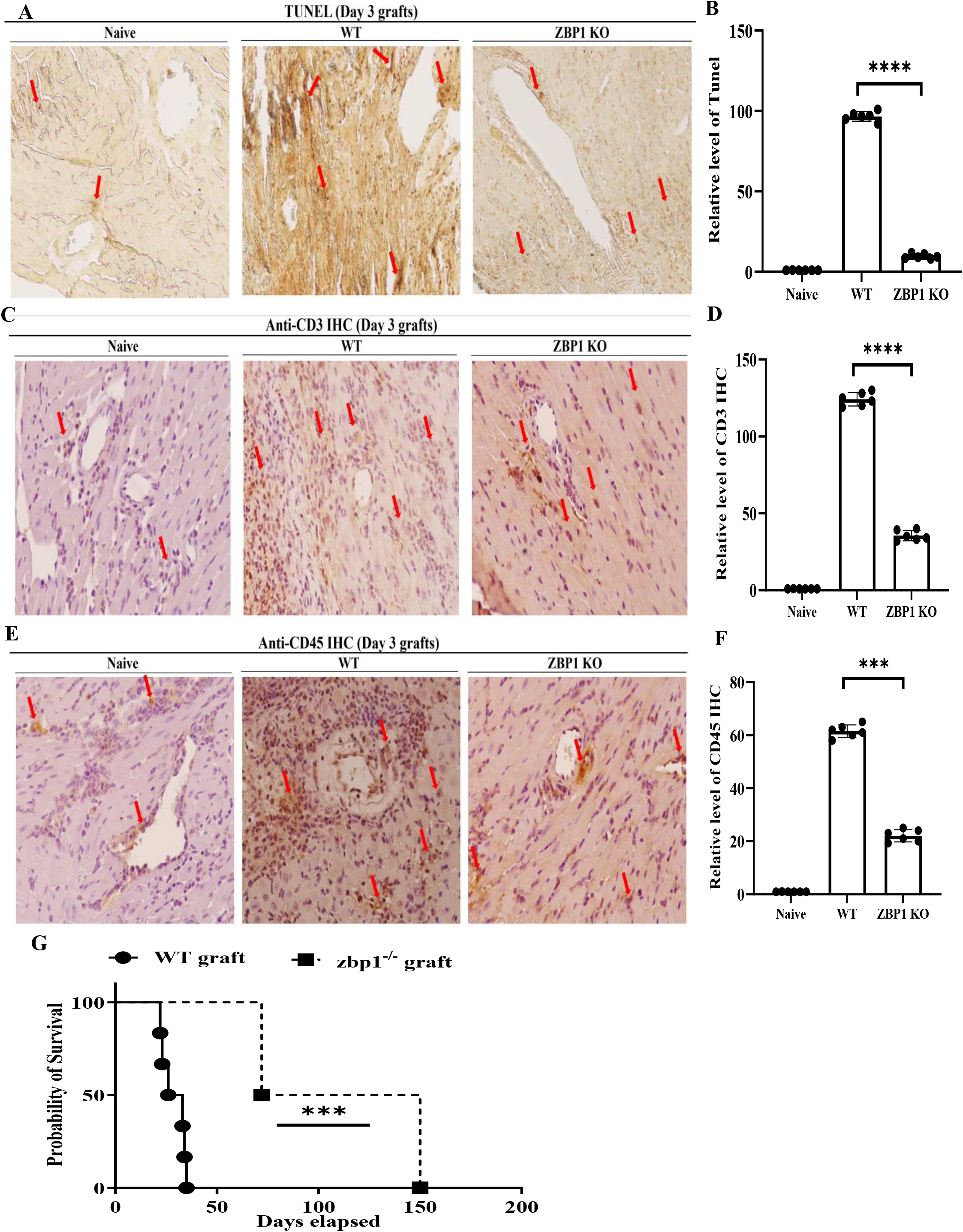
ZBP1 deficiency in the graft attenuates acute injury and induces long-term graft survival. (A&B) Cell death in heart grafts (n = 6/group) was detected by TUNEL. B6 naive hearts were used as control. Brown color indicates TUNEL-positive cells and is pointed by red arrows. Images are at 200 times magnification. Data are shown as mean± SD of 6 grafts. ****p < 0.0001, t-test. Graft lymphocyte infiltration was detected by anti-CD3 (C&D) or anti-CD45 (E&F). Red arrows indicate positive staining areas (brown color). Images were taken under 200 times magnification. Each graft was automatically counted in six connected random areas by Image J and averaged in a double-blinded fashion. **** p ≤ 0.0001. t-test. (G) Heart transplantation was performed as in Materials and Methods. All recipients received immunosuppression with anti-CD154 (0.25 mg/mouse). A cessation or significant drop in cardiac pulsation is considered graft rejection. n = 6 per group, **** p =0.00004. Log Rank test.

To determine if ZBP1 deficiency can affect transplant injury long term, the hearts from wild type B6 or ZBP1^-/-^ mice were subjected to ischemic storage as above before being transplanted into BALB/c mice. All recipients received immunosuppression with anti-CD154. Interestingly, ZBP1 deficiency benefited graft survival compared to wild-type grafts (mean survival = 128.5 ± 22.6 days vs. 28.8 ± 6.7 days, n = 6, p = 0.00014, Figure 6G).

In summary, these data demonstrate that ZBP1 deficiency attenuated graft IRI and prolonged transplant survival.

## Discussion

Transplant-associated ischemia and reperfusion and subsequent anti-donor immune responses result in cell death, tissue injury, and graft rejection. Hence, inhibition of cell death is an appropriate strategy to attenuate organ injury and prolong graft survival. However, proper cell death control has not yet been translated into clinical practice. Recent studies have revealed that multi-cell death pathways, including pyroptosis, apoptosis, and necroptosis, are activated simultaneously, known as PANoptosis. ^29^ This multi-inflammatory mechanism is driven by nuclei sensor ZBP1 and its downstream RIPK3 and caspases. We showed that ZBP1 and its ligand Z-DNA/RNA, and downstream molecules associated with PANoptosis in the graft significantly increased after transplantation. Whereas the absence of ZBP1 or the presence of ADAR1 significantly reduced cell death and graft injury and induced long-term graft survival. Our studies suggest that ZBP1 might be an important target in preventing transplant rejection.

Our study demonstrated that cfDNA contains Z-DNA and induces PANoptosis in endothelial cells (Figure 1&2), confirming that Z-DNA is the inducer of PANoptosis by stimulating ZBP1^5, 30^. Interestingly, it is reported that mitochondrial DNA (mtDNA) could be released into the cytosol and sensed by ZBP1 for cell death ^31, 32^. Although we did not measure mtDNA in cfDNA in our study, mtDNA should consist of cfDNA and stimulate ZBP1. It will be interesting to define whether mtDNA or nucleic DNA is a better inducer for ZBP1 signaling in future studies.

ADAR1 suppresses ZBP1’s function via the Zα domain interaction and thus maintains cell homeostasis. ^20, 31^ Indeed, we showed that ADAR1 could inhibit ZBP1-mediated PANoptosis. However, ADAR1 is downregulated as cell death progresses. ADAR1 downregulation or knockdown enhanced the interaction between ZBP1 and RIPK3 and thus promotes PANoptosis in endothelial cells (Figure 3). These results revealed that ADAR1 is essential for controlling the ZBP1-RIPK3 pathway. However, the mechanism for the downregulation of ADAR1 is not clear and requires further investigation. ADAR1 transgene^33^ or preventing its downregulation will be valuable strategies for preventing cell death

Nec-1s has been well reported to inhibit necroptosis by reducing the RIPK1 phosphorylation level and preventing the formation of the RIPK1-RIPK3 complex, thereby preventing necroptosis at an early stage in vitro and in vivo ^34, 35^. Hence, the release of RIPK1 from its interaction with RIPK3 might enable the transition of pRIPK3 to ZBP1. This is consistent with previous reports that RIPK1 restrain ZBP1-and TRIF-mediated cell death ^36^ or counteract ZBP1-mediated necroptosis ^37, 38^. RIPK1 not only activates RIPK3 but also nuclear factor-κB (NF-kB) in a timely manner in response to LPS or poly(I:C) ^37, 39^. We demonstrated that the level of p-NF-κB was initially increased and then decreased after 1 hour during cfDNA treatment. However, p-NF-κB stayed at a high level in the absence of RIPK3 (data not shown). Therefore, the timing and context of RIPK1-RIPK3 interaction are crucial in the early period. In addition, ZBP1 was found to protect against mtDNA-induced myocardial inflammation by activating NF-κB^32^. So, it is possible that RIPK1 can also help cell survival by activating NF-κB ^35^ or ubiquitination^40^.

In summary, our studies demonstrated that ZBP1 and its ligand Z-DNA/RNA and downstream molecules associated with PANoptosis significantly increased in the graft after transplantation. Whereas ZBP1 deficiency prevented cell death and acute graft injury and induced long-term graft survival. Hence, targeting ZBP1 could be a viable strategy to reduce transplant injury and prevent heart graft rejection.

## Acknowledgments

This study was supported by research funding from the Canadian Institutes of Health Research (CIHR, FRN169045 to ZXZ).

## Data Availability Statement

All data needed to evaluate the conclusions in the paper are present in the paper.

